# Topological morphogenesis of neuroepithelial organoids

**DOI:** 10.1101/2021.08.08.455385

**Authors:** Keisuke Ishihara, Arghyadip Mukherjee, Elena Gromberg, Jan Brugués, Elly M. Tanaka, Frank Jülicher

**Author notes:** These authors contributed equally to this work (Keisuke Ishihara, Arghyadip Mukherjee).

## Abstract

Animal organs exhibit complex topologies involving cavities and tubular networks, which underlie their form and function. However, how topology emerges during organ morphogenesis remains elusive. Here, we combine tissue reconstitution and quantitative microscopy to show that *trans* and *cis* epithelial fusion govern tissue topology and shape. These two modes of topological transitions can be regulated in neuroepithelial organoids, leading to divergent topologies. The morphological space can be captured by a single control parameter which is analogous to the reduced Gaussian rigidity of an epithelial surface. Finally, we identify a pharmacologically accessible pathway that regulates the frequency of *trans* and *cis* fusion, and demonstrate the control of organoid topology and shape. The physical principles uncovered here provide fundamental insights into the self-organization of complex tissues.

---

Tissue morphogenesis is the emergence of increasingly complex geometry and topology out of a simple group of cells (*1*). The geometry of a tissue characterises its size and shape, and its topology defines how different parts are connected and characterises the organisation of cavities and passages between them (*2–4*). Fundamental morphogenetic processes proceed as a series of size and shape changes and topological transitions (*5–8*). For example, gastrulation involves the invagination of an epithelial cell layer of initially spherical topology which eventually transitions to a toroidal topology that serves as the precursor for the passage connecting the mouth to the other end (*9*). In the case of vasculogenesis, endothelial tissues form tubular geometries that connect and result in complex topological networks with branches and loops (*10*). Fluid-filled cavities called lumens form the basis of transport networks such as bile canaliculi in the liver (*11*) and ventricles in the brain (*12*). Thus, topological and geometric transitions play a key role during morphogenesis and in organ function. However, the principles that guide the interplay of topology and geometry in formation of complex organ architectures remain unknown.

To address this fundamental issue, we make use of recent advances in 3D organ reconstitution that provide accessible and controllable experimental systems to study how a simple assembly of cells dynamically organise into tissues with complex architectures (*13, 14*). We differentiate mouse embryonic stem cells *in vitro* as free-floating aggregates that develop into neuroepithelial organoids in four days (Fig. 1a, Fig. S1a) (*15, 16*). Staining of apical surfaces (anti-ZO1, anti-PODXL) suggests that this process involves the formation of fluid-filled lumens. The apical surface of these organoids contained many passages (Fig. 1a, zoom, top panel), indicating the emergence of complex tissue geometry and topology. Here, to explore organoid morphogenesis, we set out to quantify geometry and topology and use biochemical perturbations to influence geometric and topological changes. Retinoic acid (RA) is known to act as a morphogen and to instruct the size and shape of the developing neural tube (*17*). We found that RA treatment at Day 2 of differentiation had a significant influence on the geometry and topology of organoids. Staining of apical surfaces revealed the emergence of multiple epithelial lobules by Day 4 (Fig. 1 a, +RA, zoom, bottom panel), in contrast to untreated organoids which are dominated by one large lobule (Fig. 1 a, untreated).

**Fig. 1.**
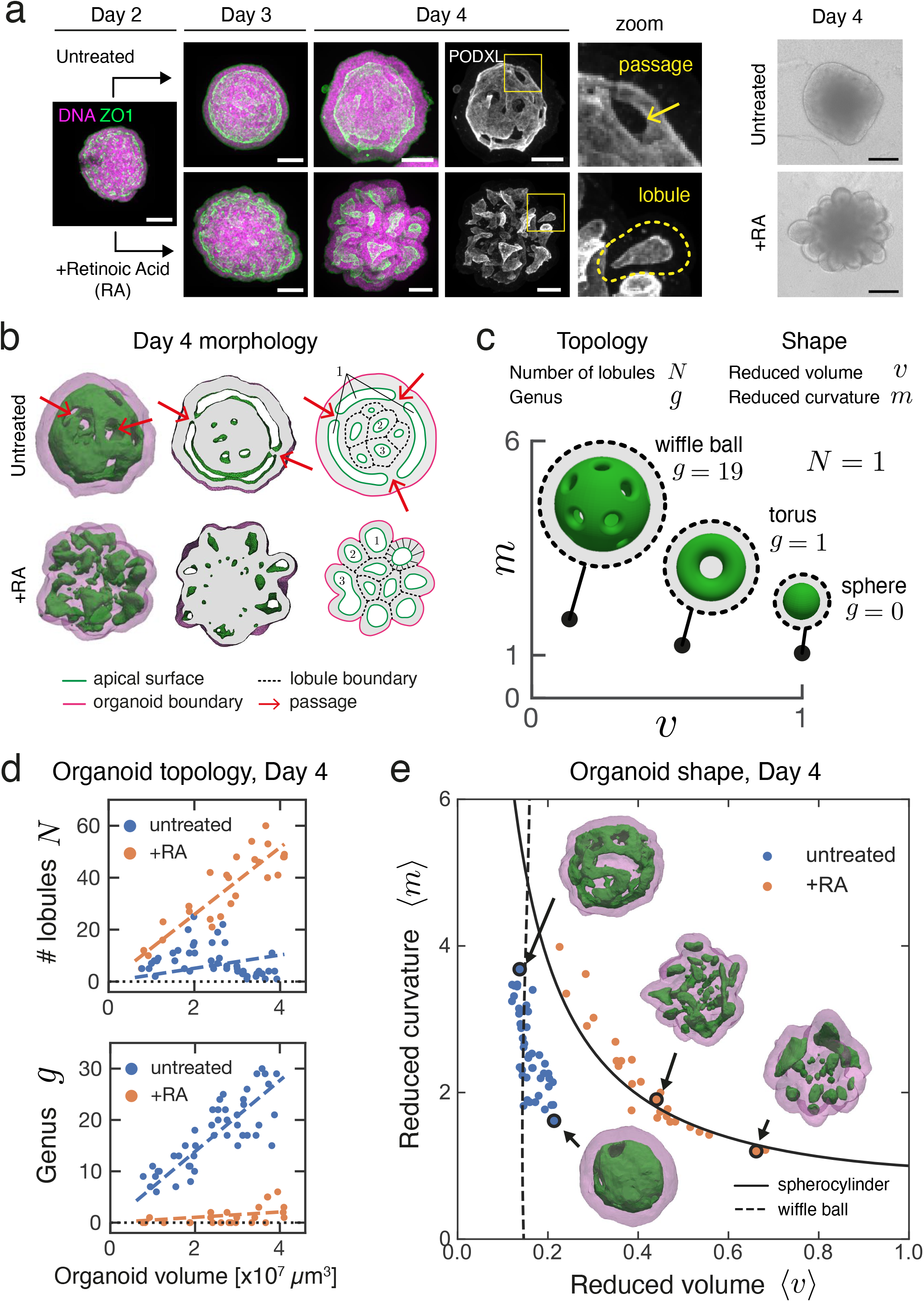
Topology and shape of neuroepithelial organoids. **a**, Neuroepithelial organoids with different morphologies were generated in the presence or absence of retinoic acid (RA). Immunofluorescence for ZO1(green) and PODXL (white) marks the apical membranes. DNA staining (magenta) marks the entire organoid. Images are maximum intensity projections of volumetric images. See Fig. S1a for images of individual confocal slices. Example of a passages on the apical surface (zoom, top row). Example of an epithelial lobule (zoom, bottom row). Brightfield images show the difference in outer morphology of organoids in the two conditions. **b**, Organoid morphology is visualised by surface renderings of organoid outer boundary (magenta, transparent) and its apical surfaces (green) for the Day 4 immunostained samples in panel a. In cross-sections, the grey regions indicate the volume occupied by cells, while the white regions indicate the fluid-filled lumens. The schematics show how organoids are divided into epithelial lobules annotated with numbers 1, 2, 3. See Fig. S1c for a schematic of a cell-level interpretation of passages. **c**, The topology of an organoid is quantified by the number of epithelilal lobules *N* and the total genus *g*. The shape of organoid is quantified by the average reduced volume ⟨*v*⟩ and average reduced curvature ⟨*m*⟩, calculated from the values of all epithelial lobules (see see Fig. S2c, Methods, and Supplementary Note) **d**, The topology of untreated (blue, n=45) and RA-treated organoids (orange, n=27) at Day 4 are characterised by the number of epithelial lobules *N* and total genus *g*, and displayed with respect to organoid volume. Organoids of different sizes were generated by varying the number of cells to be seeded at Day 0 from 300 to 2400 cells. Dashed lines are linear fits with zero intercept. **e**, The shape of untreated and RA-treated organoids are represented in the shape diagram, based on the average reduced volume ⟨*v*⟩ and average reduced curvature ⟨*m*⟩. Parametric curves for the wiffle ball (4 passages, *d/R* = 0.15, dashed line) and spherocylinders (solid line) serve as guides. Morphology of representative organoids (black circles) from both conditions are displayed. See Fig. S1e for the shapes of individual lobules in these organoids. Scale bars 100 *µm*.

In order to quantify organoid geometry and topology, we segmented the organoid architecture and constructed triangular meshes to define the outer (magenta) and apical surfaces (green) (Fig. 1b, Methods, Movie S1-S2, and Fig. S1b). This allowed us to quantify key geometric and topological measures for each organoid (Fig. S2 a,b). Using the apical surface to define distinct epithelial lobules, we characterised organoid topology by the number of lobules *N* and their topological genus *g*, and the shape of each lobule by its reduced volume *v* and reduced curvature *m* (Fig. 1c, Supplementary Note). Organoid shape is then characterised by the average of the reduced volumes and reduced curvatures over all lobules, ⟨*v*⟩ and ⟨*m*⟩, respectively (Methods, Fig. S2c). We observed that untreated organoids contained large lobules with high genus *g* that increased with organoid volume (Fig. 1d, Fig. S1d), with shape and topology that can be qualitatively captured by the wiffle ball morphology (compare Fig. 1b and c). In contrast, RA-treated organoids contained lobules of spherical topology (*g* = 0), with lobule number *N* increasing with organoid volume (Fig. 1d, Fig. S1d).

The geometry of organoids can be represented in a shape diagram (⟨*v*⟩, ⟨*m*⟩) (*18–20*)(Fig. 1e). Untreated organoids at Day 4 (blue points) had similar average reduced volumes ⟨*v*⟩ around 0.15, but varying average reduced curvature ⟨*m*⟩ falling approximately on a vertical line (dashed). This line corresponds to wiffle ball configurations for varying passage size (Fig. 1e, Supplementary Note). For RA-treated organoids at Day 4 (orange points), the average reduced curvature ⟨*v*⟩ decreases for increasing average reduced curvature ⟨*m*⟩. This trend falls on the line (solid) corresponding to spherocylinder shapes of varying aspect ratio, ranging from spheres to tubes (Fig. 1e, Supplementary Note). Thus, by Day 4, untreated organoids develop into a morphology dominated by a large lobule of high genus resembling a wiffle ball, while RA-treated organoids develop into many lobules of low topological genus consisting of spherocylinders.

To address how and when these two different organoid morphologies emerge, we imaged SiR-actin labeled organoids using light sheet microscopy over 48 hours from Day 2 to 4 (Fig. 2a, Fig. S3a, Movies S3-S4). At initial stages of imaging at Day 2, organoids in both conditions contained numerous small spherical lobules, which fused with each other and gave rise to elongated and tubular lobules (see Movies S5-S10, surface renderings). From Day 2 to 3, organoid shapes in both conditions followed a trajectory along the spherocylinder branch in the shape diagram, starting from elongated spheres (⟨*v*⟩ ≃ 1, ⟨*m*⟩ ≃ 1) and increasing in aspect ratio (Fig. 2b). At about 24 hrs of imaging (Day 3), organoid shapes reached a region in the shape diagram where the wiffle ball and spherocylinders morphologies meet (intersection of solid and dashed lines, (⟨*v*⟩ ≃ 0.15, ⟨*m*⟩ ≃ 5)). After 24 hrs, the shape trajectories for the two conditions diverged. Untreated organoids transitioned from the spherocylinder branch to the wiffle ball branch, decreasing average reduced curvature with time, while maintaining small average reduced volume (Fig. 2b, left). In contrast, RA-treated organoids remained on the spherocylinder branch, but returned towards larger average reduced volumes by Day 4 (Fig. 2b, right). During the two days of imaging, the number of lobules *N* decreased monotonically in a similar manner for both conditions (Fig. 2c, Fig. S3c). The trajectories of the total genus *g* in both conditions remained low until about 24 hrs, after which they diverged. In the untreated condition, *g* increased over time to large values from Day 3 to 4, while in the RA-treated condition, *g* remained small even at Day 4 (Fig. 2d).

**Fig. 2.**
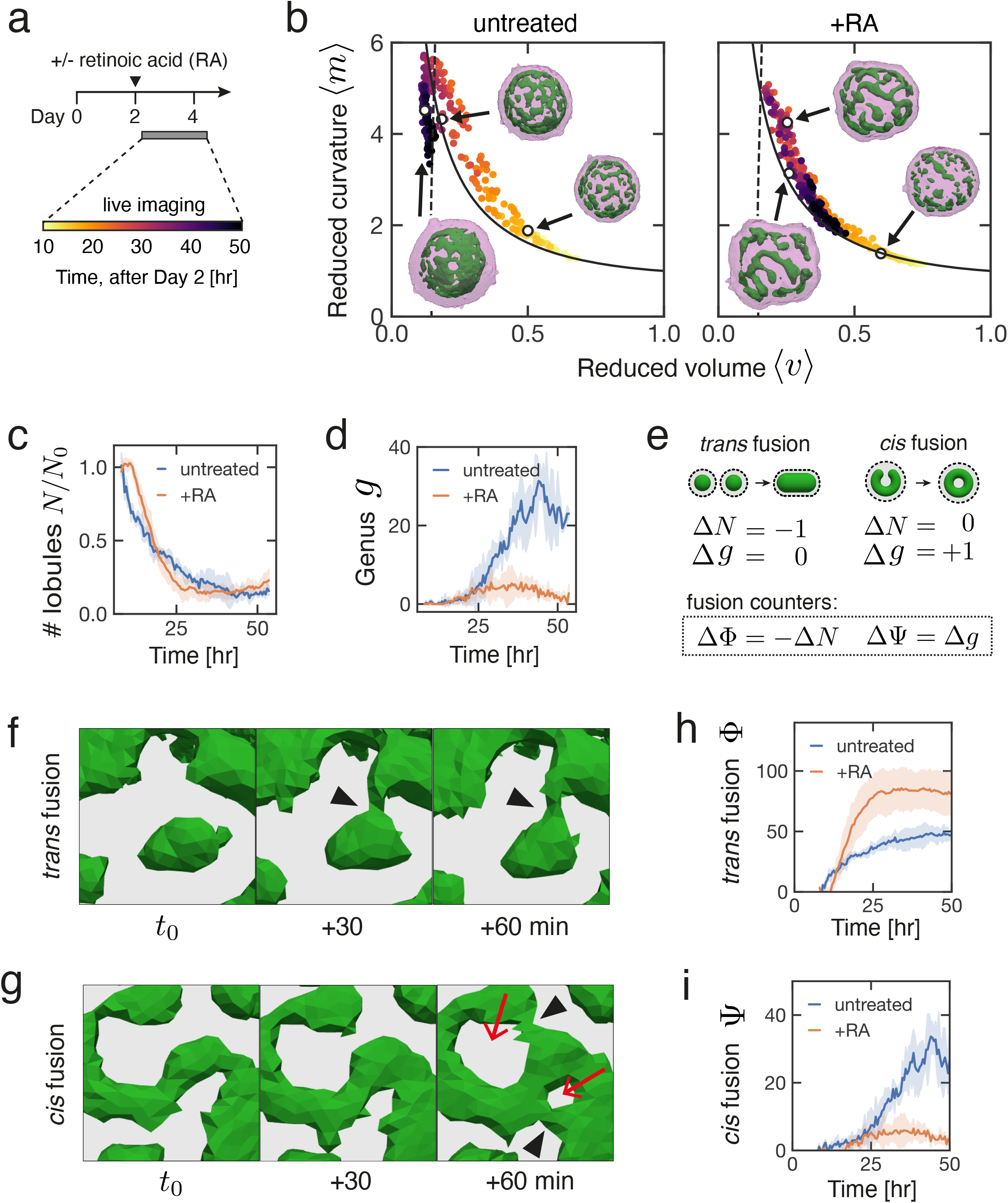
*Trans* and *cis* modes of epithelial fusion underlie emergence of organoid topology and shape. **a**, Experimental timeline for the live imaging experiment to observe organoid morphogenesis. For each time point, 3D images of SiR-actin were used to segment and reconstruct the surfaces that represent the tissue outer boundary (magenta) and apical membranes (green). **b**, Organoid shape trajectory from Day 2 to 4 for n=3 untreated organoids (left) and n=4 RA-treated organoids (right). Parametric curves for the wiffle ball (4 passages, *d/R* = 0.15, dashed line) and spherocylinders (solid line) serve as guides. Time indicates hours post RA treatment on Day 2. See Fig. S3b for shape trajectory of largest lobule in organoids. **c**, The temporal change in the number of lobules *N* normalised to the beginning of the movie for each organoid. See Fig. S3c for number of lobules *N*. **d**, The temporal change in the total genus *g* of organoids. See Fig. S3d for the temporal change in the genus of the largest lobules. **e**, *Trans* fusion and *cis* fusion are two distinct modes of topological transitions that underlie organoid morphogenesis. Counting variables for *trans* and *cis* fusion are defined based on changes in *N* and *g*. **f**, Example of *trans* fusion, in which two separate epithelial lobules fuse their apical surfaces (green). **g**, Example of *cis* fusion, in which a single epithelial lobule (apical surface, green) fuses with itself. Grey regions indicate the volume occupied by cells. Arrowheads indicate location of fusion. Red arrows indicate newly formed passages. **h**, The cumulative number of *trans* fusion events Φ over time. **i**, The cumulative number of *cis* fusion events Ψ over time. For panels c, d, h, and i, error bars indicate the standard deviation of n=3 untreated and n=4 RA-treated organoids.

The time dependence of lobule number and total genus indicated that topological transitions occur during organoid morphogenesis from Day 2 to 4. Close examination of the apical surface triangulations revealed that topological transitions occurred as two distinct modes of fusion processes. One mode of fusion, which we call *trans* fusion, involves two separate lobules that fuse with each other (Fig. 2e, left). This results in a reduction of lobule number by one (Δ*N* = −1). The second mode of fusion, which we call *cis* fusion, involves a single lobule which fuses with itself to create a passage (Fig. 2e, right). This results in an increase of genus by one (Δ*g* = +1). Experimentally observed examples of *trans* and *cis* fusion are shown in Fig. 2f and g. We defined the counters for *trans* and *cis* fusion (Fig. 2e), and quantified the cumulative fusion events from Day 2 to 4. In both conditions, the early time points are dominated by *trans* fusion events (Fig. 2h), whereas beyond the 24 hours time point, *cis* fusion is the primary mode of fusion observed only for untreated organoids (Fig. 2i). Thus, we discovered that organoid topologies emerge from distinct modes of fusion: *trans* and *cis* fusion, where only the latter leads to the creation of passages and epithelial tissues with non-spherical topology.

These findings suggest that the topology and shape of the emerging tissue are determined by the *trans* and *cis* modes of topological transitions that are generated during morphogenesis. This raises the question of which epithelial properties determine the dominant mode of fusion and how they are controlled. One possibility is that tissue mechanics (*21–23*) governs the fusion events and favors one mode of fusion over the other. An example of a mechanical property that distinguishes between the *trans* and *cis* modes of fusion is the bending energy of a fluid surface (*19, 24–27*)

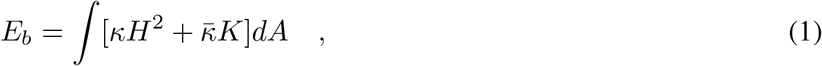

where *H* denotes the local mean curvature of the surface, *K* denotes the Gaussian curvature, and *dA* is the area element. The bending rigidity *κ* and the Gaussian rigidity 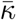 are elastic moduli that describe the resistance of the shape to bending and saddle-splay deformations, respectively. Note that ∫*KdA* = 4*π*(*N* − *g*) only depends on the indices *N* and *g* counting lobules and passages (Supplementary Note). Therefore, the Gaussian rigidity 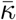 describes the resistance to topological changes. In addition, the bending rigidity *κ* governs changes in shape, but also changes in lobule number *N*. We discuss how 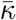 and *κ* could affect the mode of fusion by comparing the bending energy *E*_*b*_ before and after fusion (see Fig. 3a, Supplementary Note). Briefly, the change in *E*_*b*_ associated with *trans* fusion of two lobules (see Fig.3a, left) is 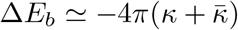; thus, *trans* fusion is energetically favoured when 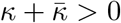. The change in energy *E*_*b*_ for *cis* fusion is 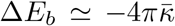 thus, passage formation via *cis* fusion in a lobule is energetically favoured when 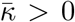 (see Fig. 3a, right). If 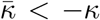, neither *trans* or *cis* fusion are energetically favoured. These three cases can be summarised in a state-diagram as a function of the ratio 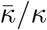 of the two elastic moduli, which is the reduced Gaussian rigidity. We can distinguish three parameter regions, where morphologies evolve differently when starting from an initial state of *N* spherical lobules (Fig. 3b). In region I, 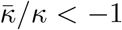, and fusion is energetically disfavoured. The system will remain in a configuration with many spherical lobules (large *N, g* = 0). In region II, 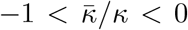, only *trans* fusion is energetically favoured. In this case, lobules will tend to fuse, resulting in fewer lobules of spherical topology (small *N, g* = 0). In region III, 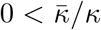, both *trans* and *cis* fusion are energetically favoured. As a result, large lobules with many passages exemplified by wiffle ball morphology emerge (small *N*, large *g*). Our results suggest that RA treatment shifts the behavior of the system from region III to region II. To test this idea, we estimate the scaled bending energy *E*_*b*_*/κ* of lobule geometries as a function of time considering different values of reduced Gaussian rigidity 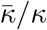 (see Methods, Fig. 3c, left, and Fig. S4). Only for positive 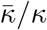 corresponding to region III, scaled bending energy *E*_*b*_*/κ* decreases with time for untreated organoids, consistent with the idea that untreated organoids operate in region III. After RA treatment, a decrease in scaled bending energy *E*_*b*_*/κ* with time requires a shift of 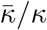 to negative values (Fig. 3c, right). This suggests that after RA-treatment the mechanical properties of the epithelia is modified, possibly leading to a reduction of 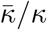 below zero at later times.

**Fig. 3.**
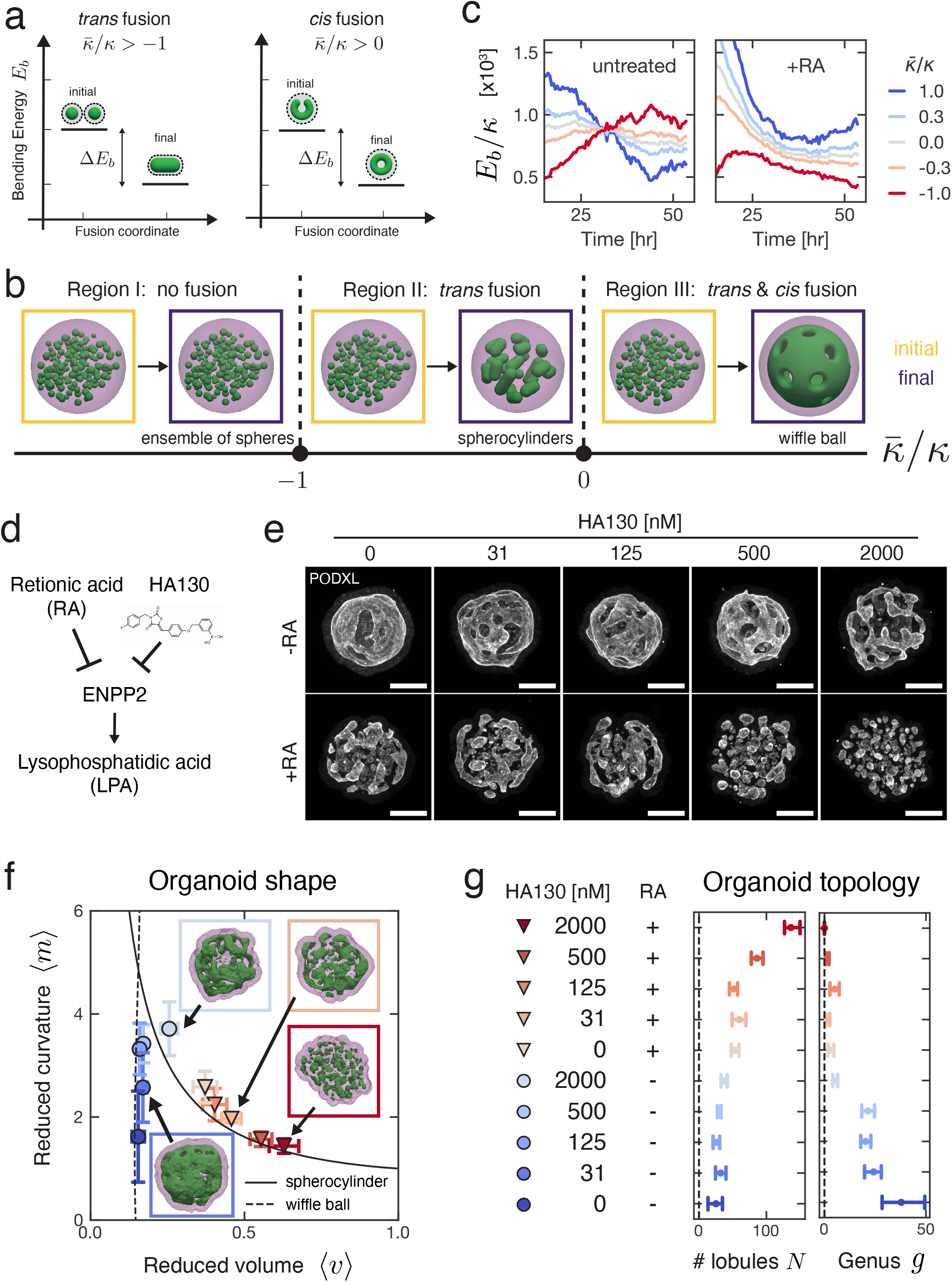
Energetics of *trans* and *cis* fusion and pharmacological control of organoid morphogenesis. **a**, Change in bending energy Δ*E*_*b*_ is depicted for *trans* and *cis* fusion. The conditions for Δ*E*_*b*_ *<* 0 is denoted for both fusion modes in terms of topological rigidity 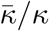. **b**, Temporal evolution of scaled bending energy is shown for various values of 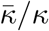, estimated from triangulated meshes 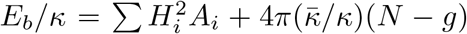 where the indices *i* traverses all faces in the system. See Fig. S4a for the evolution of individual terms and Fig. S4b for the corresponding energy for the largest lobule in the system. Estimations were made for the same n=3 untreated and n=4 RA-treated organoids represented in Fig. 2. **c**, A state-diagram of organoid morphology is shown as a function of reduced Gaussian rigidity 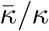. Three regions depict initial (yellow) and final (purple) morphologies: Region I, 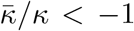: an ensemble of spheres is stable and no fusion is favoured. Region II, 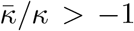 *trans* fusion leads to formation of tubular lobules with spherical topology. Region III, 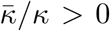 both *trans* and *cis* fusion is favoured and leads to lobules with wiffle-ball like morphology. **d**, Retinoic acid treatment of organoids induces the transcriptional down-regulation of ENPP2, which is responsible for lysophosphatidic acid (LPA) synthesis. HA130 is a pharmacological inhibitor for ENPP2 (*31*). **e**, HA130 treatment leads to organoid morphologies that are consistent with attenuation of epithelial fusion, both in the absence (top row) and presence (bottom row) of RA. See Movie S11 and S12 for surface rendering examples. **f**, Shape of organoids cultured in varying concentrations of HA130 (see panel **g** for legend). Organoid morphology is visualised by surface renderings of organoid outer boundary (magenta, transparent) and its apical surfaces (green) for the Day 4 samples shown in panel **e**. Parametric curves for the wiffle ball (4 passages, *d/R* = 0.15, dashed line) and spherocylinders (solid line) serve as guides for the shape diagram. **g**, Topology of organoids cultured in varying concentrations of HA130. Errorbar indicates 95% confidence interval from n=5-10 organoids per condition.

Thus, the primary effect of RA-treatment on topological morphogenesis is consistent with a decrease of the reduced Gaussian rigidity 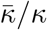. This raises the question whether we can control tissue mechanical properties corresponding to 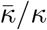 by molecular perturbations. To identify such molecular pathway, we performed RNAseq on organoids treated with or without RA (see Methods, Table S1, and Fig. S5). We found that RA treatment down-regulates the expression of ENPP2, which is the cellular enzyme that produces lysophosphatidic acid (LPA) (*28*) (Fig. 3d). LPA is known to induce the enlargement of apical membrane area of human neuroepithelial cells (*29*) and enhance the folding of developing mouse brains (*30*). To test the role of LPA synthesis on organoid morphogenesis, we treated Day 2 organoids with varying concentrations of HA130, a small molecule inhibitor of LPA synthesis (*31*) (Fig. 3e), and analysed their morphology at Day 4 (Fig. 3f, top and Fig. S6a, b). At low concentrations of HA130 (31-125 nM), organoids had primarily a single lobule with many passages, resembling wiffle ball morphologies, similar to the untreated case and consistent with region III (Fig. 3f, g). Increasing the concentration of HA130 results in morphologies with fewer passages until at the highest concentration of HA130 used (2 *µ*M), organoids resembled a set of elongated spheres, consistent with the gradual shift from region III to region II (Fig. 3e, top right, Movie S11). This perturbation shows that inhibition of LPA synthesis mimics the effect of RA-treatment in organoid morphogenesis. Our theory based on reduced Gaussian rigidity predicts that further reduction of 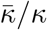 can result in a transition from region II to region I, where both *trans* and *cis* fusion are disfavoured. This behavior was not observed in the presence or absence of RA. To test this prediction, we applied varying concentrations of HA130 to RA-treated organoids, thus combining the effect of RA-treatment and inhibition of LPA synthesis (Fig. 3e, bottom and Fig. S6a, c). For increasing concentrations of HA130 (31-2000 nM), RA-treated organoids exhibited increasing numbers of lobules (Fig. 3g) consistent with gradual suppression of *trans* fusion. At the highest concentrations of HA130 (500 and 2000 nM), RA-treated organoids resulted in an ensemble of spherical lobules with lobule number *N* at Day 4 comparable to the Day 2 organoids (Fig. 3e bottom right, Movie S12). This is consistent with the suppression of fusion and a shift to region I in the state diagram (Fig. 3b). By combining the effects of RA-treatment with HA130, we achieved a shift of organoid morphogenesis from region III via region II to region I. The corresponding organoid shapes followed a trajectory in the (⟨*v*⟩, ⟨*m*⟩) diagram starting from the wiffle ball branch (Fig. 3f, dashed) at low (*m*), moving upwards when shifting towards region II. The trajectory transitions to the spherocylinder branch (Fig. 3f, solid) when passages are no longer observed corresponding to region II. The trajectory follows this branch towards large (*v*) and small (*m*) as the system shifts to region I. The pharmacological modulation of LPA synthesis together with RA treatment allowed us to tune topological morphogenesis by controlling the modes of fusion and their rates, giving rise to a broad range of morphologies with varying shape and topology.

In summary, by combining quantitative 4D microscopy, theory, and pharmacological perturbations, we have discovered how topological transitions drive neuroepithelial morphogenesis through *trans* and *cis* epithelial fusion. The morphological space can be captured by a single control parameter which is analogous to the reduced Gaussian rigidity 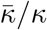 of an epithelial surface. We envision that the physical principle of topological morphogenesis uncovered here could apply to other biological structures from sub-cellular organelles to complex tissues.

## Supporting information

Supplementary Information including methods

## Acknowledgements

The authors thank the members of Brugués, Tanaka, and Jülicher groups for discussions. We thank Marta Shahbazi for providing mouse ES cell lines and discussions. We thank the Light Microscopy Facility and Technology Development Studio at MPI-CBG, in particular, Rico Barsacchi and Marc Bickle for assistance with the automated confocal imaging. We thank DRESDEN-concept Genome Center at TU Dresden and MPI-CBG for assistance with RNAseq analysis. We thank the BioOptics and Bioinformatics facilities at IMP and IMBA for assistance with imaging and RNAseq analysis.

## Author contributions

This project was conceived by KI, AM, JB, EMT, and FJ. KI designed and performed all the experiments except for the RNAseq experiment performed by EG. Results were analyzed by KI and AM. Theoretical models were conceived by KI, AM, JB, and FJ. Main text was written by KI, AM, JB, EMT, and FJ.

## Data and materials availability

All materials are available upon request.

## Supplementary Materials

- Figs. S1-S6
- Materials and Methods
- Supplementary Note
- Movie S1-S12

