## Supplementary Information including methods for "Topological morphogenesis of neuroepithelial organoids"

### Supplementary Information
### Topological morphogenesis of neuroepithelial organoids

Keisuke Ishihara[1,2,3,4,5,†], Arghyadip Mukherjee[1,2,3,4,†], Elena Gromberg[5],
Jan Brugués[1,2,3,4,*], Elly M. Tanaka[5,*], Frank Jülicher[2,3,4,*]

[1]Max Planck Institute of Molecular Cell Biology and Genetics (MPI-CBG); 01307 Dresden, Germany.

[2]Max Planck Institute for the Physics of Complex Systems (MPI-PKS); 01187 Dresden, Germany.

[3]Center for Systems Biology Dresden; 01307 Dresden, Germany.

[4]Cluster of Excellence Physics of Life, TU Dresden; 01062 Dresden, Germany.

[5]Research Institute of Molecular Pathology (IMP), Campus-Vienna-Biocenter 1, 1030 Vienna, Austria.

[†]These authors contributed equally to this work.

**Table of contents:**

- Fig S1-S6
- Materials and Methods
- Supplementary Note
- Caption for Table S1
- Captions for Movies S1-S12

Figure S1

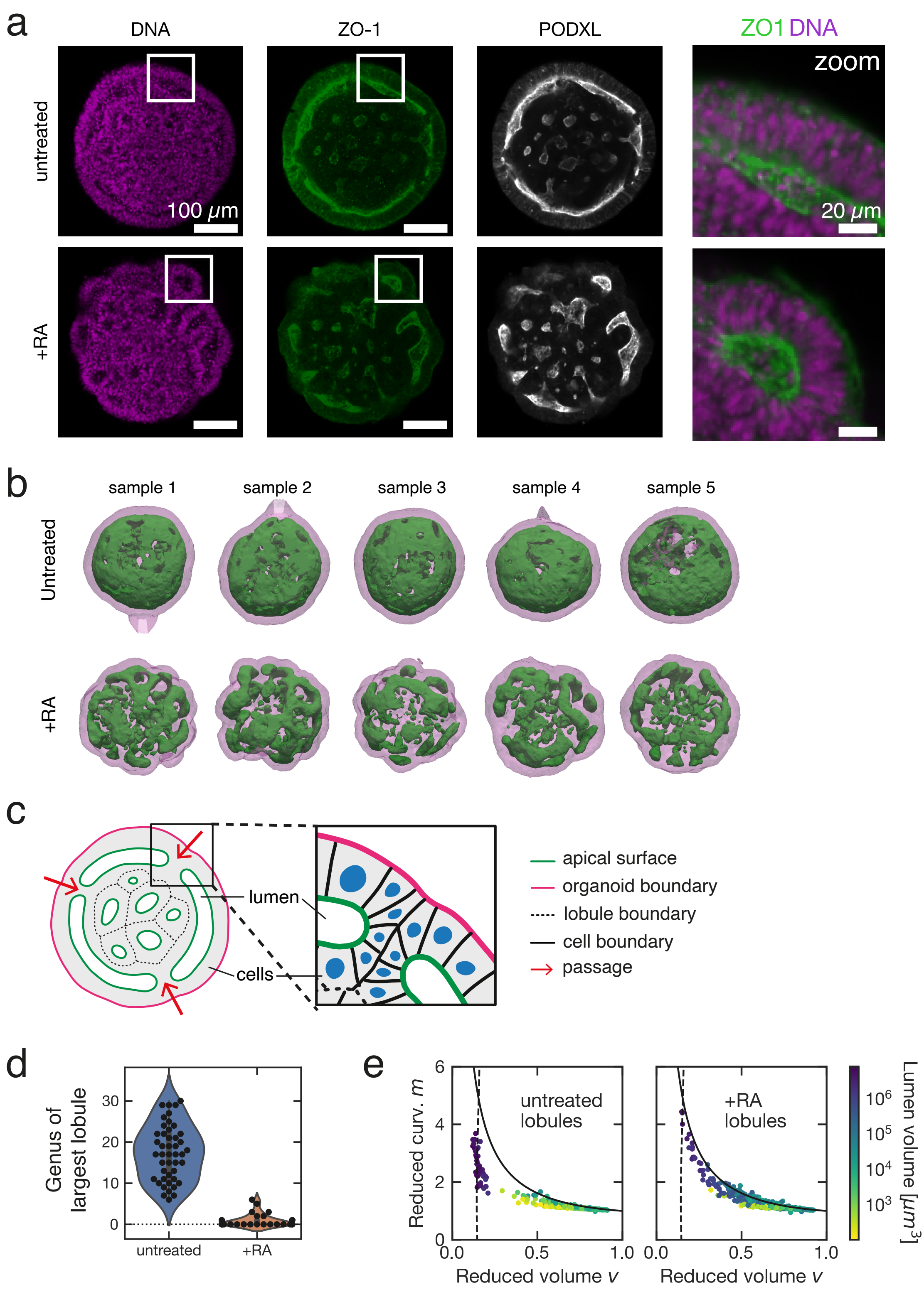

a
DNA
ZO-1
PODXL
ZO1 DNA
zoom
untreated
+RA
100 μm
20 μm
b
sample 1
sample 2
sample 3
sample 4
sample 5
Untreated
+RA
c
lumen
cells
apical surface
organoid boundary
lobule boundary
cell boundary
passage
d
Genus of largest lobule
untreated
+RA
e
Reduced curv. m
Reduced volume v
untreated lobules
+RA lobules
Lumen volume [μm³]

**Figure S1 a,** Single planes from spinning disc confocal microscopy images of optically cleared organoids stained for DNA, anti-ZO1, and anti-PODXL. **b,** Examples of untreated and RA-treated organoids at Day 4. Surface representations show organoid outer boundary (magenta, transparent) and apical surfaces (green). **c,** Schematic representation of an organoid cross-section, describing the relation between the organoid outer boundary (magenta), apical surfaces (green), lobule boundary (dotted lines), cell nuclei (blue), and passages (red arrow). **d,** Genus of the largest lobule in each organoid. **e,** Shape diagram of individual lobules found in multiple organoids. Lumen volume is indicated by colour. Solid line: spherocylinders. Dashed line: wiffle ball. Panels **b** through **c** show data from the same n=45 untreated and n=27 RA-treated organoids as in Fig. 1.

### Figure S2

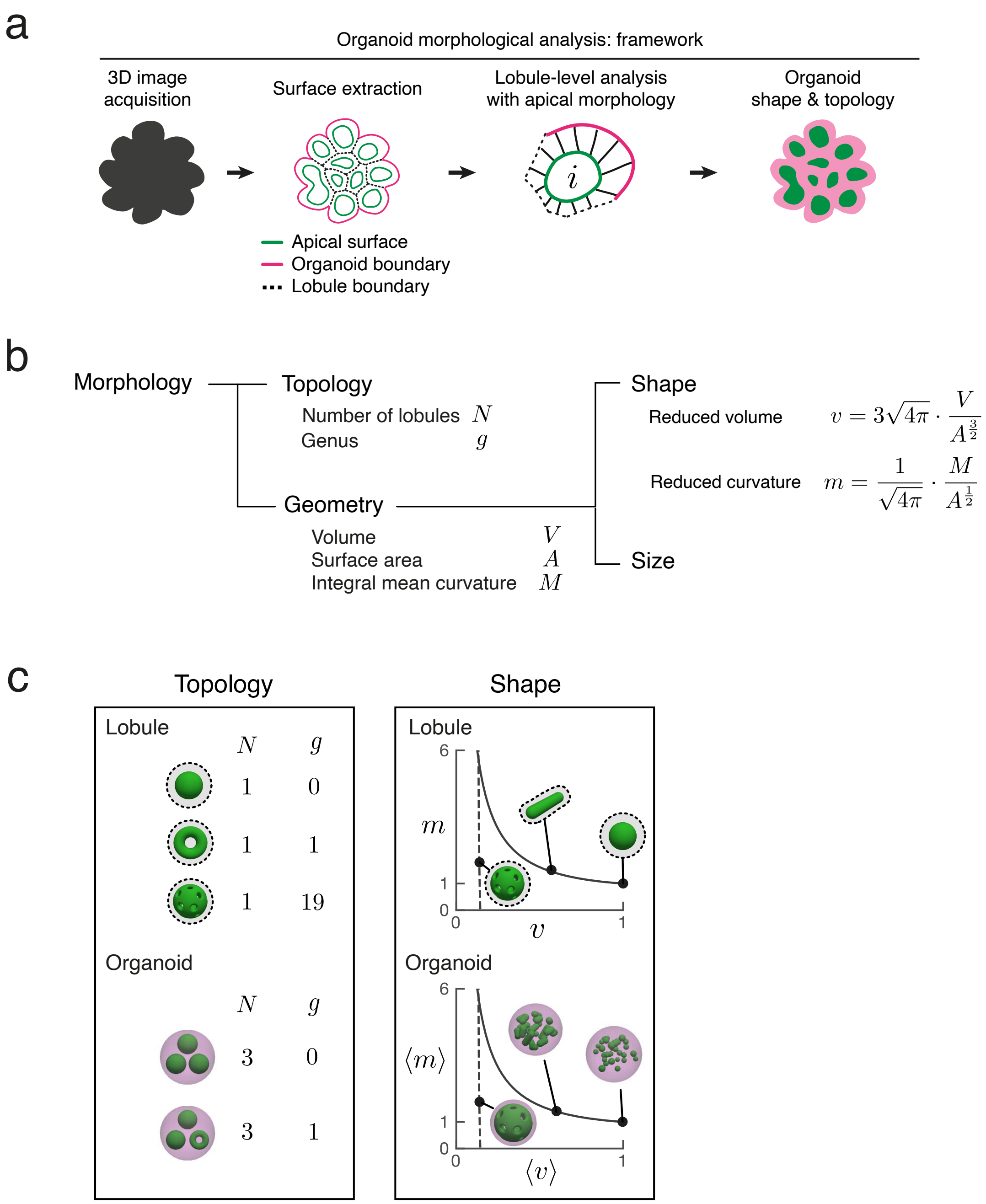
a
Organoid morphological analysis: framework
3D image acquisition
Surface extraction
Lobule-level analysis with apical morphology
Organoid shape & topology
i
Apical surface
Organoid boundary
Lobule boundary
b
Morphology
Topology
Number of lobules $N$
Genus $g$
Geometry
Volume $V$
Surface area $A$
Integral mean curvature $M$
Shape
Reduced volume $v = 3\sqrt{4\pi} \cdot \frac{V}{A^{\frac{3}{2}}}$
Reduced curvature $m = \frac{1}{\sqrt{4\pi}} \cdot \frac{M}{A^{\frac{1}{2}}}$
Size
c
Topology
Lobule
$N$ $g$
1 0
1 1
1 19
Organoid
$N$ $g$
3 0
3 1
Shape
Lobule
6
$m$
1
0
0
$v$
1
Organoid
6
$\langle m \rangle$
1
0
0
$\langle v \rangle$
1

**Figure S2 a,** Framework for morphological analysis of organoids. First, we acquire 3D images of a tissue sample, here, an organoid. Second, we perform image segmentation and surface construction of triangulated meshes to represent 3D tissue morphology as a set of apical surfaces and organoid outer boundary. Next, we use individual apical surfaces to define epithelial lobules and characterize their morphology. Finally, we integrate the information from all lobules to define the overall topology and shape of an organoids. This process is repeated for multiple organoids. **b,** In mathematical terms, morphology is discussed for the aspects of topology and geometry. Topology describes the connectivity of objects, which do not change from stretching or shrinking. Geometry is discussed in terms of the shape and size of objects. Shape is a scale invariant quality, while size has the units related to length scale. **c,** The metrics for topology are the number of lobules $N$ and total genus $g$ in an organoid. The metrics for shape are the reduced volume $v$ and reduced curvature $m$ of individual lobules, which are averaged to yield organoid level quantities $\langle v \rangle$ and $\langle m \rangle$ (see Methods).

### Figure S3

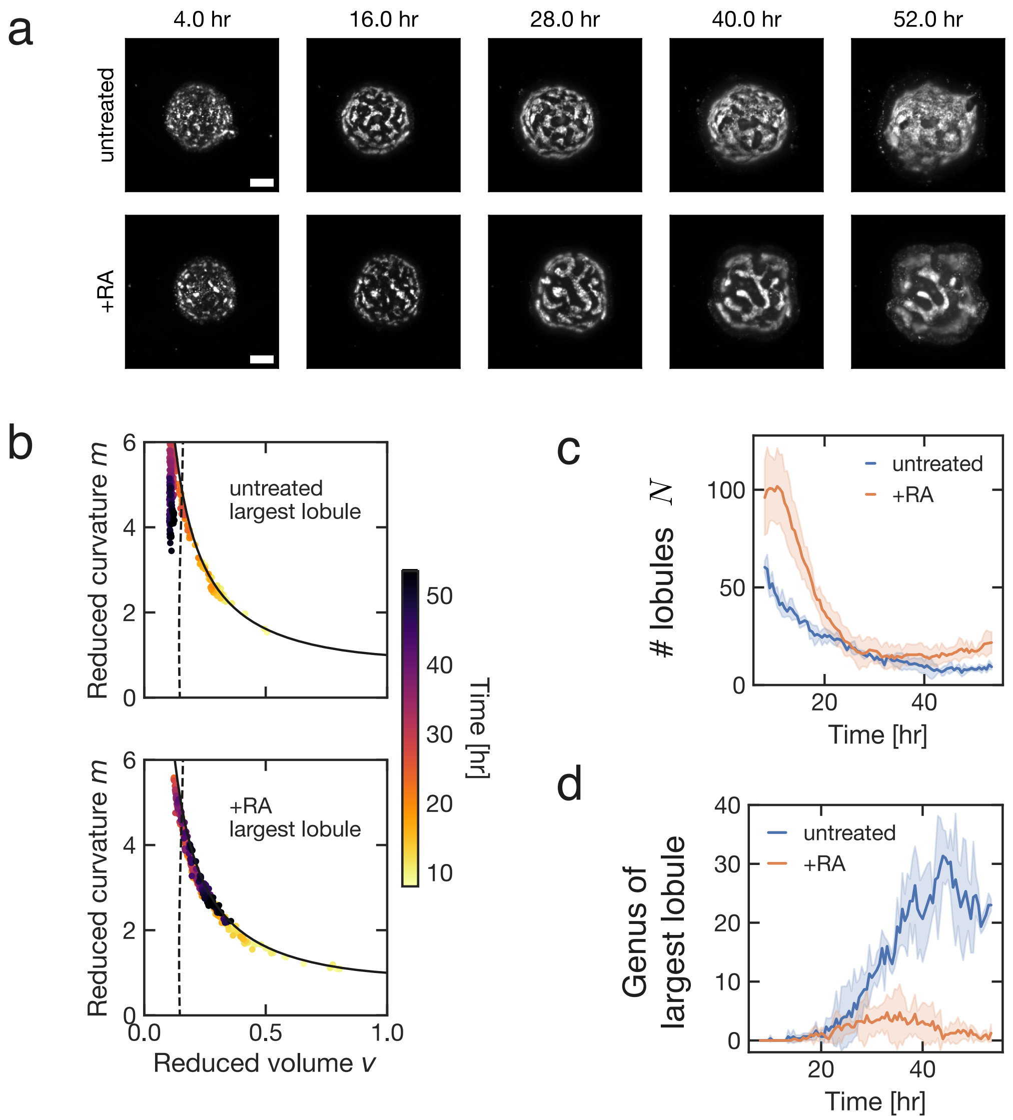
a
4.0 hr
16.0 hr
28.0 hr
40.0 hr
52.0 hr
untreated
+RA
b
untreated
largest lobule
+RA
largest lobule
Reduced curvature m
Reduced volume v
Time [hr]
c
### lobules N
untreated
+RA
Time [hr]
d
Genus of
largest lobule
untreated
+RA
Time [hr]

**Figure S3 a,** Individual frames from light sheet imaging experiment of organoids. Images shown are maximum intensity projections of SiR-actin signal. Time indicates time post RA treatment. **b,** Shape diagram displays how the shape of the largest lumen in the organoid evolves over time. Solid line: spherocylinder. Dashed line: wiffle ball. **c,** The temporal change in the number of lobules $N$ in organoids. **d,** The temporal change in the genus $g$ of the largest lobule of each organoid. Panels show data from the same n=3 untreated and n=4 RA-treated organoids as in Fig. 2. Scale bar, 100 $\mu m$.

### Figure S4

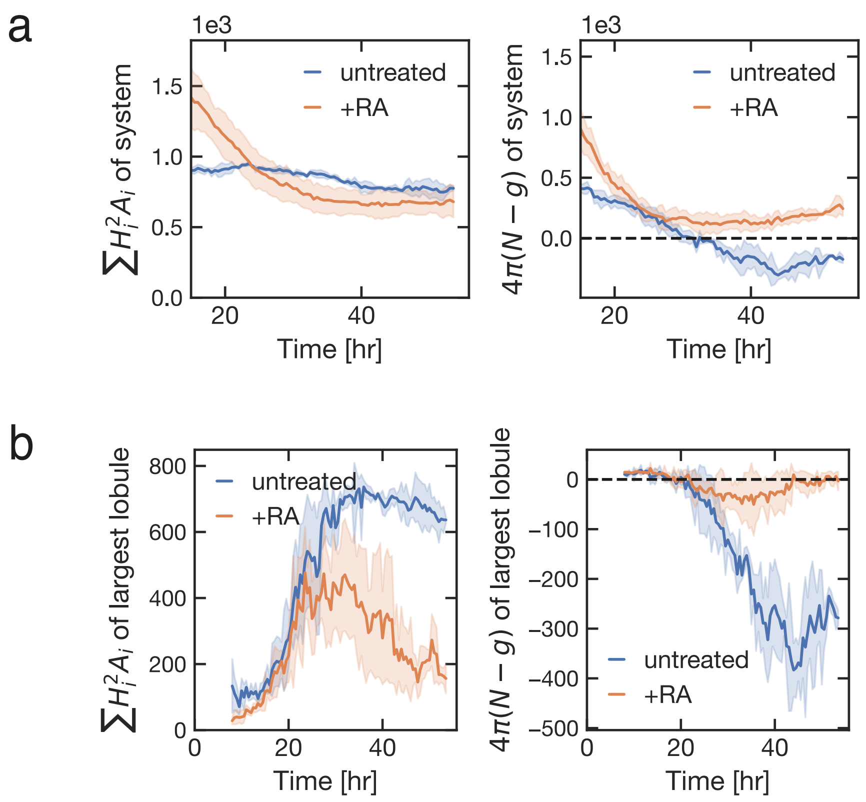
a
1e3
1.5
1.0
0.5
0.0
20
40
Time [hr]
untreated
+RA
$\sum H_i^2 A_i$ of system
1e3
1.5
1.0
0.5
0.0
20
40
Time [hr]
untreated
+RA
$4\pi(N-g)$ of system
b
800
600
400
200
0
0
20
40
Time [hr]
untreated
+RA
$\sum H_i^2 A_i$ of largest lobule
0
−100
−200
−300
−400
−500
0
20
40
Time [hr]
untreated
+RA
$4\pi(N-g)$ of largest lobule

**Figure S4 a,** Temporal evolution of bending elasticity ($\sum_i H_i^2 A_i$) and Gaussian elasticity ($\int K dA = 4\pi(N-g)$) is depicted for organoids. **b,** Temporal evolution of bending elasticity ($\sum_i H_i^2 A_i$) and Gaussian elasticity ($\int K dA = 4\pi(N-g)$) is depicted for largest epithelial lobules of each organoids. Errorbars indicate the standard deviation of the same n=3 untreated and n=4 RA-treated organoids represented in Fig. 2.

Figure S5

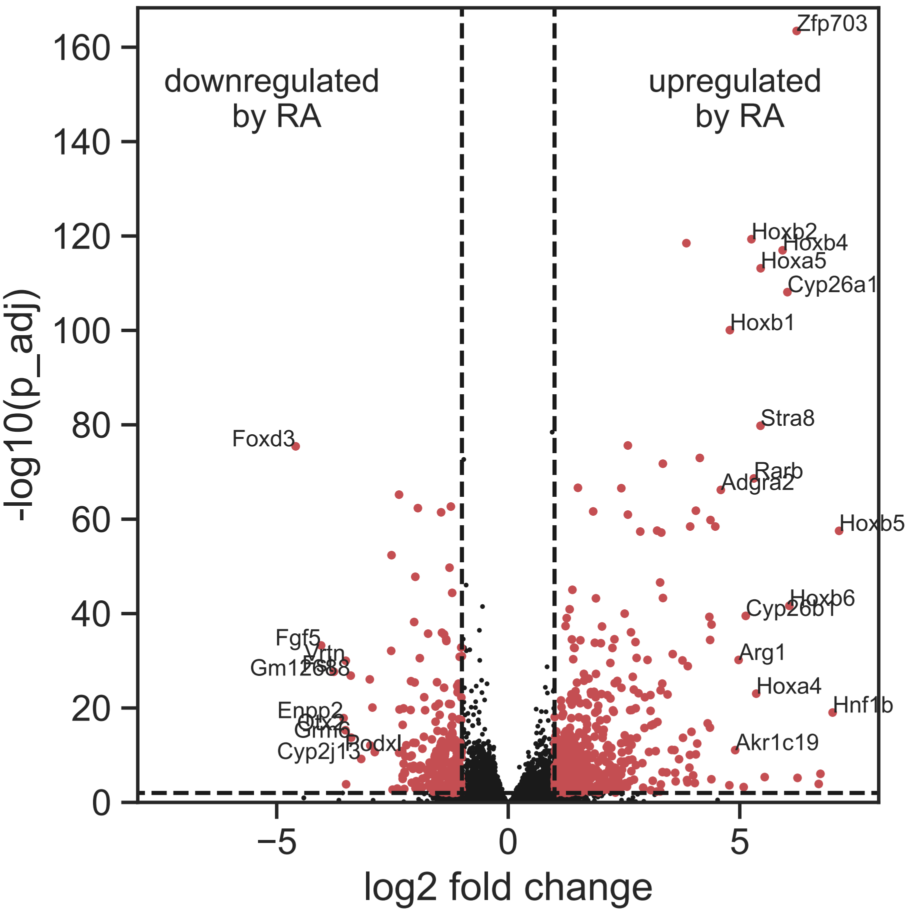
downregulated
by RA
upregulated
by RA
-log10(p_adj)
log2 fold change
Zfp703
Hoxb2
Hoxb4
Hoxa5
Cyp26a1
Hoxb1
Stra8
Rarb
Adgra2
Hoxb5
Hoxb6
Cyp26b1
Arg1
Hoxa4
Hnf1b
Akr1c19
Foxd3
Fgf5
Vrtn
Gm12688
Enpp2
Otx2
Grm6
Podxl
Cyp2j13

**Figure S5** Volcano plot displaying gene expression change in organoids treated with or without retinoic acid (RA). Red points represent all differentially regulated genes (see Methods and Table S1 for a complete list).

Figure S6

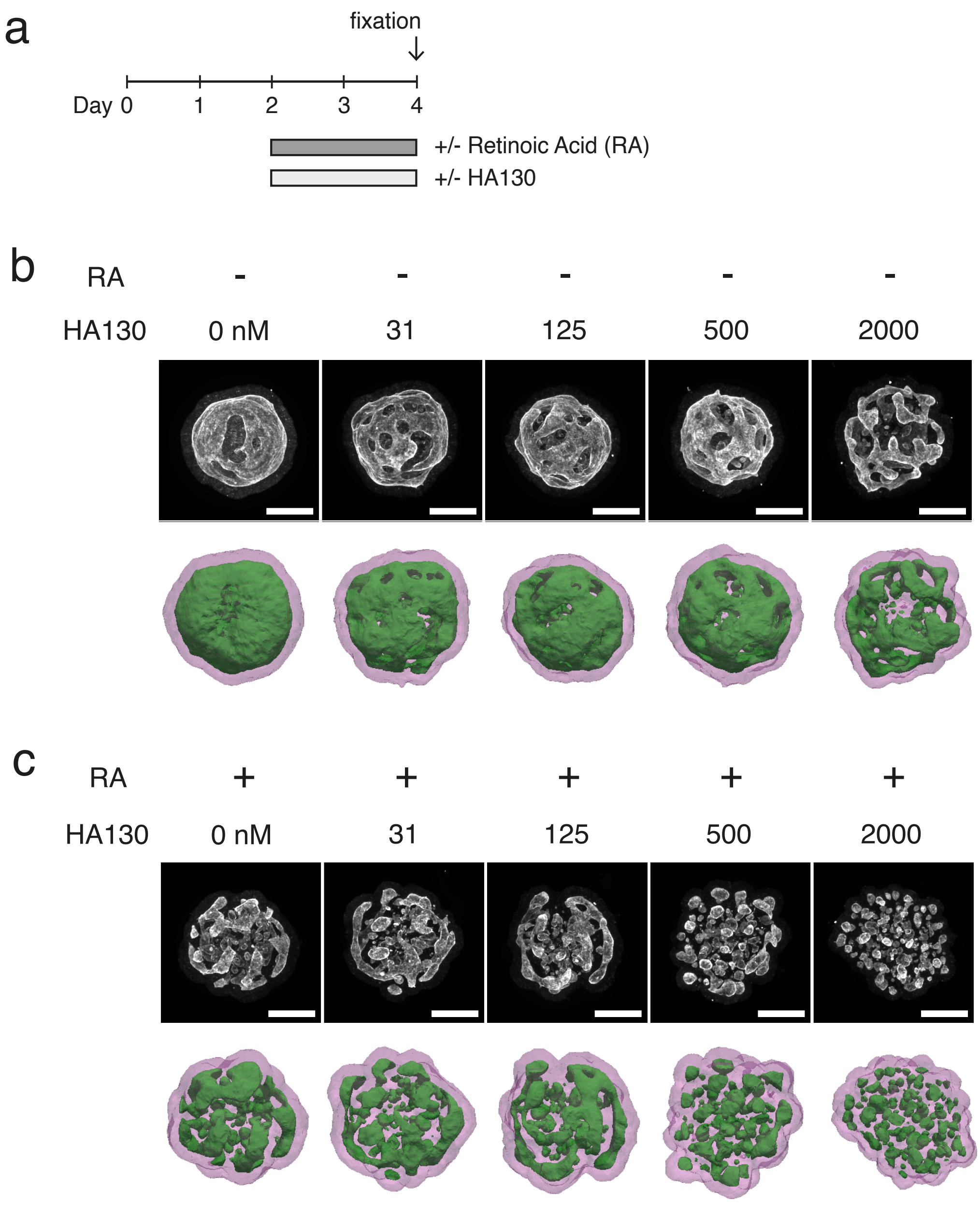
a
fixation
Day 0
1
2
3
4
+/- Retinoic Acid (RA)
+/- HA130
b
RA
-
-
-
-
-
HA130
0 nM
31
125
500
2000
c
RA
+
+
+
+
+
HA130
0 nM
31
125
500
2000

**Figure S6 a,** Workflow of experiment to test the role of lysophosphatidic acid (LPA) synthesis in organoid morphogenesis. Organoids were treated with varying concentrations of HA130, a small molecular inhibitor for ENPP2, with or without 250 nM retinoic acid (RA) during Day 2 to Day 4. Organoid morphologies were scored at Day 4. **b,** Representative examples of Day 4 organoids treated with varying concentrations of HA130 in the absence of RA. **c,** Representative examples of Day 4 organoids treated with varying concentrations of HA130 in the presence of RA. Images show maximum intensity projections of PODXL immunofluorescence signal. Organoid outer boundary (magenta) and apical surfaces (green) are shown for the corresponding sample. Scale bar, 100 $\mu$m.

### Materials and Methods

#### Mouse embryonic stem cell and neuroepithelial organoid culture.

Mouse embryonic stem (ES) cells were passaged in N2B27 medium with 2i/LIF following standard protocols (*1*). To generate neuroepithelial organoids, we modified a published protocol for optic cup organoids (*2*). On Day 0, ES cells were dissociated into single cells and re-aggregated in N2B27 neural induction medium (typically 1000 cells per 50 $\mu$l ml per well) in 96-well U-bottom low adhesion plates (ThermoFisher, 174925). On Day 1, Matrigel (BD Biosciences, 354234) dissolved in 50 ul N2B27 medium was added to each well for a final concentration of 2.5% (v/v). On Day 2, 50 $\mu$l N2B27 with or without all-trans retinoic acid (RA, Sigma R2625) was added to each well. The final concentration of RA was 250 nM. In some experiments, organoids were treated with varying concentrations of HA130 (Echelon Biosciences) to inhibit lysophosphatidic acid synthesis. All 3D morphological data in this study was collected from neuroepithelial organoids made from E14 mouse ES cells (a gift from Marta Shahbazi, MRC Laboratory of Molecular Biology). The major morphological phenotypes induced by the addition of retinoic acid and HA130 were confirmed by experiments using R1 wild-type mouse ES cells (a gift from Ronald Naumann, MPI-CBG).

#### Immunofluorescence and optical clearing of orgnaoids.

Organoid samples were washed in PBS, fixed with ice cold 4% paraformaldehyde solution for 30 min, rinsed, and stored in PBS at 4C. Organoids were permeabilised and blocked in blocking solution (PBS + 5% (w/v) Bovine Serum Albumin + 0.3% (v/v) Triton X-100) for 1 hr at room temperature. Primary antibodies were diluted in 50 $\mu$l blocking solution and applied to organoids overnight with occasional mixing. After removal of primary antibody solution, organoids were washed with 0.5 ml PBST (PBS + 0.3% Triton X-100), 2 hr x 3 rounds with occasional mixing. Secondary antibodies and DNA stain were diluted in 50 $\mu$l blocking solution overnight with occasional mixing. After removal of secondary antibody solution, organoids were washed with 0.5 ml PBST, 2 hr x 3 rounds and finally stored in PBS. Primary antibodies used were: Rat anti-PODXL (R&D, MAB1556) at 1:40 dilution, Mouse anti-ZO1 (Invitrogen 33-9100) at 1:40 dilution. Secondary antibodies used were: Alexa568 anti-rat at 1:100 dilution, Alexa647 anti-mouse at 1:100 dilution. DNA stain was SytoxGreen (S7020, Invitrogen) used at 84 nM. Immunostained organoid samples were optically cleared using the 2nd generation ethyl cinnamate clearing protocol (*3*).

#### Fixed sample imaging and 3D segmentation.

Optically cleared organoids were transferred to an ethylcinnamate-resistant 96 well plate (Ibidi, #89621) for high throughput imaging. Imaging was performed with an automated spinning disk microscope (Yokogawa, CellVoyager 7000S) equipped with a CSU-W1 spinning disk and a CMOS camera (1280x1080 pixels). To cover an entire well from a 96 well plate, four fields of view were acquired with a 4x objective lens. These overview images were processed on the fly via a custom ImageJ macro executed by the SearchFirst module in the Wako Software Suite which identified the coordinates of individual organoids. These coordinates were subsequently revisited with a 20x air (NA = 0.75) Olympus objective. For each position, 300 planes with 0.8 $\mu$m spacing were acquired. Due to the 'fish-bowl' effect of ethyl cinnamate's high refractive index (RI=1.56), the effective z-steps were 1.25 $\mu$m and the entire z-stack encompassed 374 $\mu$m in sample depth. We used 2x2 binning on the camera which resulted in images at 0.648 $\mu$m/pixel. For image analysis, we further binned the images in xy so that the 3D voxel dimensions were comparable (1.30 x 1.30 x 1.25 $\mu$m in x, y, and z). To segment the organoid outer boundary, we applied Otsu thresholding to the z-stacks from the SytoxGreen channel. To segment the apical surface, we applied Otsu thresholding to the z-stacks from the PODXL channel. All reported measurements have accounted for the effect of tissue shrinkage (factor of 0.603 in linear dimensions, experimentally determined) of the ethylcinnamate clearing protocol.

#### Live imaging and 3D segmentation.

Neuroepithelial organoids were cultured as described above with the addition of 100 nM SiR-actin (Spirochrome) in the medium from Day 1. Immediately after RA treatment on Day 2, organoids were transferred to custom multiwell chambers and imaged on a light sheet microscope (Viventis, LS1) equipped with a sCMOS camera and 638 nm laser line. Every 30 min, z-stacks were acquired at 3 micron intervals, covering a sample depth of 200 microns. For image analysis, we binned the images in xy so the voxel dimensions were nearly isotropic (2.76 x 2.76 x 3.00 $\mu$m in x, y, and z). To segment the organoid outer boundary, we applied multi-Otsu thresholding (classes=3 using lowest threshold) to the SiR-actin z-stacks. To segment the apical surface, we applied multi-Otsu thresholding (classes=4 using highest threshold) to the SiR-actin z-stacks.

#### Surface construction and morphological analysis.

The marching cubes algorithm was used to extract triangulated meshes from the segmented 3D images that represent the organoid outer boundary and apical surfaces. To reduce mesh complexity, a combination of PyMesh functions were used to collapse short edges (tolerance=2x the minimum voxel length of the input image), remove duplicated vertices, remove duplicated faces, and remove degenerated/obtuse triangles. For each closed surface, we calculated the volume $V$, surface area $A$, integral mean curvature $M$, and Euler characteristic $\chi$ using functions implemented in PyMesh and our custom code (see Supplementary Note). These quantities were used to calculate the reduced volume $v = 3\sqrt{4\pi}\ V/A^{3/2}$, reduced curvature $m = M/\sqrt{4\pi A}$, and the topological genus $g = 1 - \chi/2$ of epithelial lobules. Epithelial lobules with lumen volume $V < 100\ \mu\text{m}^3$ were excluded from analysis. Organoid level quantities were defined as total quantities $g = \sum_i g_i$ or as weighted averages $\langle v \rangle = \sum_i v_i V_i / \sum_i V_i$ and $\langle m \rangle = \sum_i m_i V_i / \sum_i V_i$ where the index $i$ enumerates all $N$ epithelial lobules of an organoid. All computational analysis and visualization in this study were performed using Python 3.7 with the libraries NumPy (*4*), SciPy (*5*), Pandas (*6*), Matplotlib (*7*), Seaborn (*8*), Scikit-Image (*9*), PyMesh (*10*), PyVista (*11*), and Polyscope (*12*).

#### RNA sequencing and data processing

Organoids were grown in 35 mm MatTek dishes by seeding 3000-5000 mouse ES cells in 50 $\mu$l Matrigel and cultured in 2 ml N2B27 medium at 37C, 5% CO2 as described in (*13, 14*). At Day 2, organoids were treated with or without 250 nM all-trans retinoic acid. Organoids were extracted from the Matrigel with Cell Recovery solution (Corning 354253) at 0, 18, 30, 42, 54, 68, and 80 hrs post Day 2. Three biological replicates were prepared for each cell population, yielding 39 independent samples for RNA extraction. For each sample, a minimum of 50k cells were collected for RNA extraction using the QIAGEN RNeasy Micro Kit. RNAseq libraries were prepared using the NEBNext Poly(A) mRNA Magnetic Isolation Module (E7490) and the NEBNext Ultra Directional RNA Library Prep Kit for Illumina (E7420) according to the manufacturer's instructions. For ligation custom adaptors were used (Adaptor-Oligo 1: 5'-ACA CTC TTT CCC TAC ACG ACG CTC TTC CGA TCT-3', Adaptor-Oligo 2: 5'-P-GAT CGG AAG AGC ACA CGT CTG AAC TCC AGT CAC-3'). The libraries were sequenced 75bp single end on an Illumina NextSeq500 system. RNA-seq reads were trimmed using trimgalore v0.5.0, filtered to remove abundant sequences using bowtie2 v2.3.4.1, aligned to the GRCm38 genome (Ensembl release 94) using star v2.6.0c and summarized per gene with featureCounts (subread v1.6.2). Further analysis was performed using DESeq2 v1.18.1. To identify significantly differentially regulated genes between untreated versus RA-treated cell populations, we focused on the 18 and 30 hrs post Day 2 time points, and selected genes with $\log_2$ fold change less than -1 or greater than 1 and false discovery rate of 0.01 (based on adjusted p-values) in both time points, identifying 569 up-regulated and 250 down-regulated genes.

### Supplementary Note

#### 1 Morphological analysis of epithelial lobules and organoids

Here we provide a brief summary of geometric concepts used in the analysis of epithelial lobules, and an extended discussion on the geometric theory of epithelial surfaces.

##### 1.1 Integral geometry of smooth surfaces

We consider the apical surface of the epithelial lobules as closed smooth surfaces. At any point on the surface, the curvature is captured by the mean curvature

$$H = \frac{1}{2}\left(C_1 + C_2\right) \tag{1}$$

and the Gaussian curvature

$$K = C_1 C_2 \quad , \tag{2}$$

where $C_1, C_2$ are the principal curvatures (Fig. SN1a).

Integral geometry provides a set of "global" morphological descriptors called Minkowski functionals that characterize the geometry and topology of objects (*15*). In 3-dimensional space, they are

$$\text{Volume:} \quad V = \int dV \tag{3}$$

$$\text{Surface Area:} \quad A = \int dA \tag{4}$$

$$\text{Integral Mean Curvature:} \quad M = \int H dA \tag{5}$$

$$\text{Euler characteristic:} \quad \chi = \frac{1}{2\pi}\int K dA = 2(N-g) \quad , \tag{6}$$

where $dV$ and $dA$ are the volume and area element, respectively. Topology is captured by the Euler characteristic $\chi$ of the object, which relates to the Gaussian curvature via the Gauss-Bonnet theorem $2\pi\chi = \int K dA$. For a single object, $\chi = 2 - 2g$ where $g$ is the topological genus that quantifies the number of handles. For an epithelial organoid consisting of multiple epithelial lobules, $\chi = 2(N - g)$ where $N$ is the number of distinct lobules and $g$ is total genus.

##### 1.2 Integral geometry of discretized surfaces

Volumetric images of organoids are segmented and used to construct triangular meshes to represent the apical surfaces of epithelial lobules. Thus, we use discrete approximations of the Minkowski functionals Eq.(3-6) to characterize geometry and topology. The meshes are defined as polyhedron, with $N_v$ number of vertices connected via $N_e$ number of edges, and encapsulate $N_f$ number of flat faces. The surface area $A$ and volume $V$ then are calculated as

$$A = \sum_i^{N_f} A_i \quad , \tag{7}$$

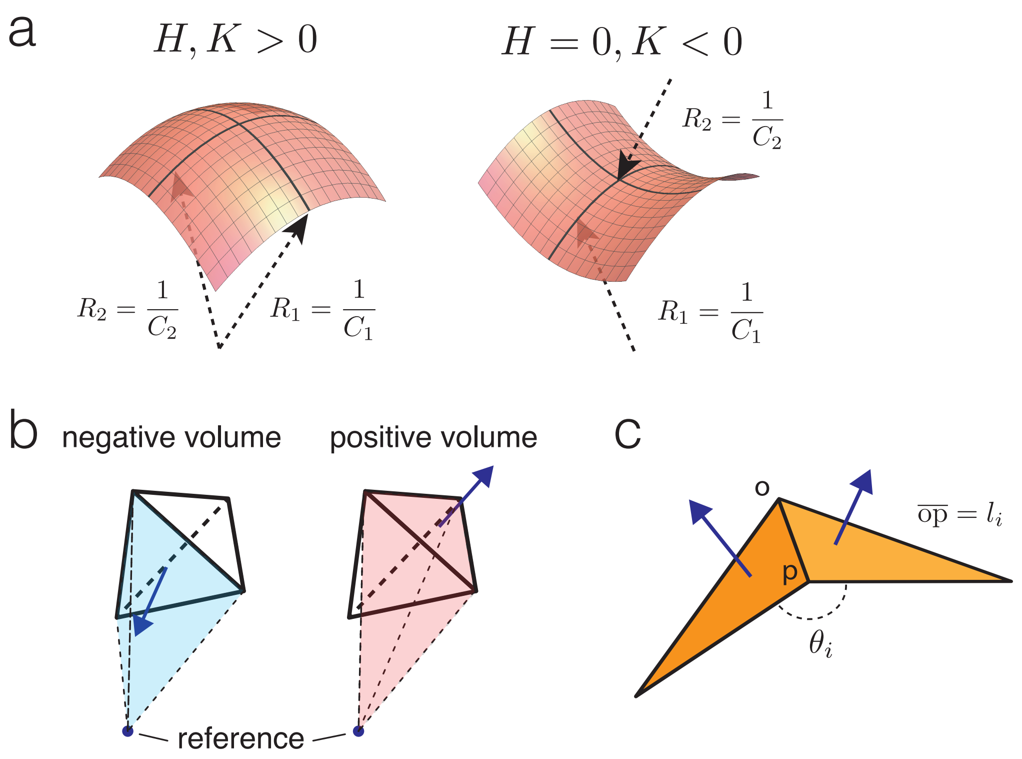


Fig. SN1: **a**, Principle curvatures $C_1$ and $C_2$ and associated radii of curvature $R_1$ and $R_2$ are shown for a convex (left) and saddle (right) surface. Mean curvature $H$ and Gaussian curvature $K$ are indicated. **b**, Signed volumes (negative or positive) are indicated for depicted tetrahedrons based on the position of the reference point. **c**, Dihedral angle $\theta_i$ between two adjacent faces is shown, the associated edge $\overline{\text{op}}$ has a length of $l_i$.

and

$$V = \sum_{i}^{N_f} V_{\uparrow,i} \quad , \tag{8}$$

where $A_i$ is the face area of the i$^{th}$ face and $V_{\uparrow,i}$ is the signed volume of a tetrahedron defined by the three vertices of the face and an arbitrary reference point (Fig. SN1b). The sign of $V_{\uparrow,i}$ is determined by asking if the face normal vector is pointing towards (negative volume) or away (positive volume) from the reference point.

For the integral mean curvature $M$, we follow Steiner's approach (*16*) to mollify the polyhedron, or smoothen its edges and vertices, with a ball of radius $\epsilon > 0$. In the limit of very small $\epsilon$, we obtain

$$M = \sum_{i}^{N_e} \frac{1}{2}\theta_i l_i \quad , \tag{9}$$

where $l_i$ is the edge length and $\theta_i$ is the dihedral angle at the edge ( (*17*), Fig. SN1c).

The Euler characteristic $\chi$ of the polyhedron is calculated using Euler's formula,

$$\chi = N_v - N_e + N_f \quad . \tag{10}$$

##### 1.3 Morphological analysis with shape diagrams

We use two non-dimensional metrics, reduced volume $v$ and reduced curvature $m$ to capture the shapes of epithelial lobules,

$$\text{Reduced volume:} \quad v \equiv 3\sqrt{4\pi}\frac{V}{A^{3/2}} \tag{11}$$

$$\text{Reduced curvature:} \quad m \equiv \frac{M}{\sqrt{4\pi A}} \quad . \tag{12}$$

$v$ is an isoperimetric quantity that captures the compactness of a surface in three-dimensions, while $m$ measures the curvature-area mismatch. For a sphere, both $v$ and $m$ are equal to unity. In the following section, we analyze idealized shapes of spherocylinders and wiffle balls with shape diagrams $(v, m)$.

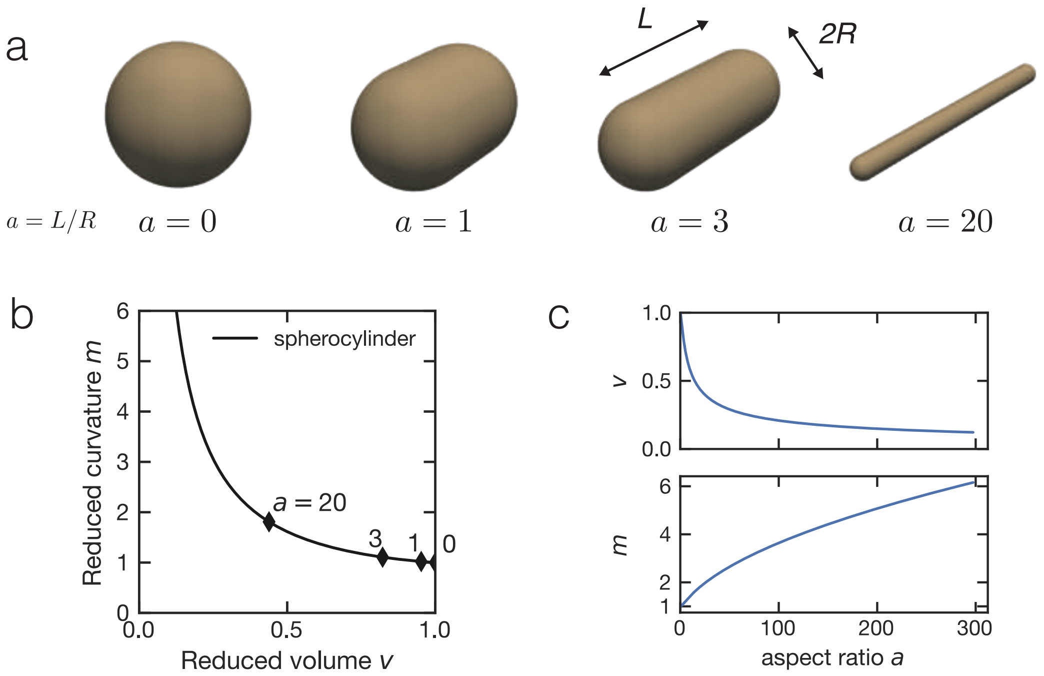


Fig. SN2: **a**, Spherocylinders of increasing (left to right) aspect ratio $a$ are shown. **b**, Spherocylinders with specific values of $a$ are represented as points in the shape diagram. The solid-line represents the parametric curve for the spherocylinder family. **c**, Reduce volume $v$ (top) and reduced curvature $m$ (bottom) are shown as a function of aspect ratio $a$.

###### 1.3.1 Spherocylinder

During morphogenesis, neuroepithelial lobules start from spherical shapes and become increasingly elongated tubular structures. The spherocylinder provides an idealised model to investigate this experimentally observed shape transition. A spherocylinder consists of a cylinder of length $L$ with spherical caps at both ends, where the radius of the cylinder and of the spheres are both $R$ (Fig. SN2a). This shape can be parametrized by the aspect ratio $a = L/R$. For aspect ratio $a = 0$, we recover a sphere. For higher values of aspect ratio $a$, the spherocylinder represents increasingly elongated tubes. The Minkowski functionals are then given by,

$$\begin{aligned} V =& \quad \pi R^2 L + \frac{4\pi}{3} R^3 = \pi R^3 (a + \frac{4}{3}) && (13) \\ A =& \quad 2\pi R L + 4\pi R^2 = 2\pi R^2 (2 + a) && (14) \\ M =& \quad \frac{1}{2R} \cdot 2\pi R L + \frac{1}{R} \cdot 4\pi R^2 = \pi R (a + 4) && (15) \\ \chi =& \quad 2 && (16) \end{aligned}$$

The reduced volume $v$ and reduced curvature $m$ can be parametrized with the aspect ratio $a$ as,

$$v = \frac{3a/4 + 1}{(1 + a/2)^{3/2}}, \tag{17}$$

$$m = \frac{a/4 + 1}{\sqrt{1 + a/2}} \quad . \tag{18}$$

These equations define a characteristic curve for spherocylinder in the shape diagram (Fig. SN2b,c).

##### 1.3.2 Wiffle ball

The final geometry of the Day 4 neuroepithelial lobules resemble a wiffle ball, a familiar children's toy in North America (Fig. SN3b). We consider two concentric spheres with radii $R$ and $R + d$, connected via $p$ number of passages (see Fig. SN3,a-b). For simplicity we consider all identical passages with central aperture $2\theta$ (Fig. SN3c). We model the passages with the inner surface of a torus with major radius $r = (R + d/2)\sin\theta$ and minor radius $d/2$.

We obtain $\theta_{\max}(p)$ using approximate solutions for the optimal packing of $p$ points on a spherical surface (*18*). To a good approximation $\theta_{\max}(p) \sim \sqrt{\pi/p}$ . The minima of the aperture angle $\theta_{\min}(d/R)$ is set by the condition $r > d/2$.

Note that for a wiffle ball with $p$ passages the genus $g = p - 1$. The Minkowski functionals are given by,

$$\begin{aligned} V =& \quad \frac{4\pi - p\,\Omega_p}{3} \left( (R + d)^3 - R^3 \right) + p\,V_p && (19) \\ A =& \quad (4\pi - p\,\Omega_p) \left( (R + d)^2 + R^2 \right) + p\,A_p && (20) \\ M =& \quad (4\pi - p\,\Omega_p) d + p M_p && (21) \\ \chi =& \quad 2 - 2g = 2 - 2(p - 1) = 4 - 2p \quad . && (22) \end{aligned}$$

Here $V_p$, $A_p$, $M_p$ and $\Omega_p$ are respectively the volume, surface area, integral mean curvature, and the solid angle of one passage,

$$\begin{aligned} V_p =& \quad \frac{\pi}{2} \left( \frac{d}{2} \right)^2 \cdot 2\pi \left( r - \frac{4}{3\pi} \frac{d}{2} \cos\theta \right) = \frac{\pi^2}{4} d^2 r - \frac{\pi}{6} d^3 \cos\theta && (23) \\ A_p =& \quad \pi d \int_{\theta - \pi}^{\theta} d\theta' \left( r + \frac{d}{2} \sin\theta' \right) = \pi^2 d r - \pi d^2 \cos\theta && (24) \\ M_p =& \quad \pi d \int_{\theta - \pi}^{\theta} d\theta' \left( r + \frac{d}{2} \sin\theta' \right) \left( \frac{2}{d} + \frac{\sin\theta'}{r + \frac{d}{2}\sin\theta'} \right) = 2\pi^2 r - 4\pi d \cos\theta && (25) \\ \Omega_p =& \quad \int d\Omega = \int_0^{\theta} d\theta' 2\pi \sin\theta' d\theta' = 2\pi(1 - \cos\theta) \quad . && (26) \end{aligned}$$

We find that for $d/R << 1$, to a leading order $v \sim d/R$. The variation in reduced curvature $m$ arises from dependencies on $\theta$ and $p$, which to a linear order can be approximated to,

$$m \sim \sqrt{\frac{\pi^3}{2}}\theta p - \sqrt{2\pi}\,\frac{d}{R}(p-1) \quad . \tag{27}$$

As a result, the parametric curve for a wiffle ball in the shape diagram is almost a vertical line (Fig. SN3d, e).

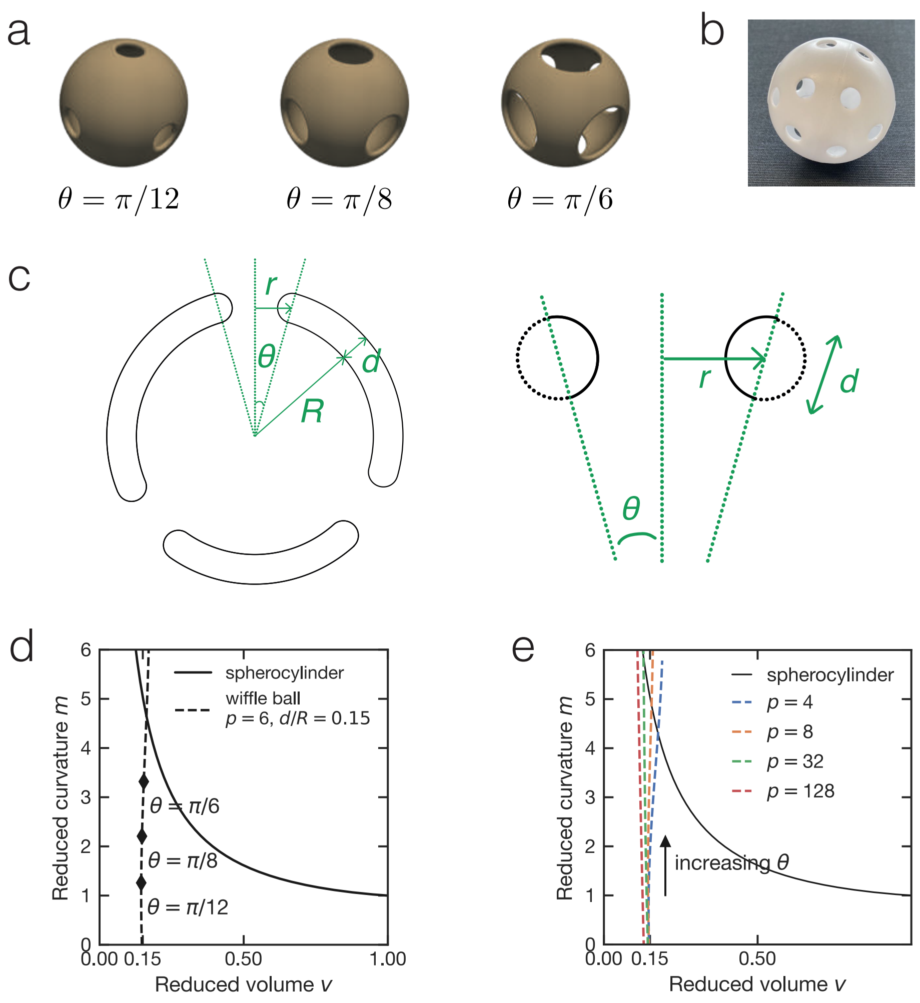


Fig. SN3: **a**, Wiffle balls with $p = 6$ passages are shown for varying opening angle $\theta$. **b**, Image showing a wiffle ball, a familiar children's toy in North America. **c**, Left, cross-sectional diagram of a wiffle ball depicting geometric variables; thickness $d$, inner radius $R$ and opening angle $\theta$. Right, cross-sectional view of a passage of the wiffle ball; geometric variables are indicated. **d**, Parametric curve (dashed line) for a wiffle ball with $p = 6$ passages is shown for $d/R = 0.15$. **e**, Parametric curve (colored dashed lines) for a wiffle ball with varying number of passages (see legend) is shown with $d/R = 0.15$.

### 2 Topological transitions in a system of fluid surfaces

In this section we discuss topological transitions in a system of fluid surfaces with $N$ closed surfaces or lobules and $g$ number of handles. The transition in topology of the system is thus characterized by changes in the topological indices $N$ and $g$. The mechanics of such fluid surfaces can be captured with a bending energy

$$E_b = \int \left(\kappa H^2 + \bar{\kappa} K\right) dA \quad , \tag{28}$$

where $H$ , $K$ an $dA$ respectively denote the local mean curvature, the Gaussian curvature of the surface, and the area element. The bending rigidity $\kappa$ and the Gaussian rigidity $\bar{\kappa}$ are elastic moduli that capture the resistance of the shape to bending and saddle-splay deformations, respectively. Due to Gauss-Bonnet theorem, $\int K dA = 2\pi\chi = 4\pi(N - g)$ is a topological invariant and depends only on the number of surfaces $N$ and total genus $g$. The Gaussian rigidity $\bar{\kappa}$ hence describes the resistance to topological changes that occur via changes in $N$ as well as $g$. The bending rigidity $\kappa$ dictates changes in shape, but also changes in $N$.

#### 2.1 Energetics of topological transitions

For a sphere with radii $R$, the mean curvature $H = 1/R$, Gaussian curvature $K = 1/R^2$ and the bending energy is given by,

$$\begin{aligned} E_b &= \int \left(\kappa H^2 + \bar{\kappa} K\right) dA \\ &= \left(4\pi R^2 \frac{1}{R^2}\kappa + 4\pi R^2 \frac{1}{R^2}\bar{\kappa}\right) \\ &= 4\pi(\kappa + \bar{\kappa}) \quad . \end{aligned} \tag{29}$$

Note that $E_b$ is independent of $R$ and hence only captures aspects of shape. For a system of $N$ spheres of any size $E_b = 4\pi N(\kappa + \bar{\kappa})$. In such a system a decrease in $N$ reduces $E_b$ iff $\kappa + \bar{\kappa} > 0$. Change in bending energy for a *trans* fusion is $\Delta E_b - \simeq 4\pi(\kappa + \bar{\kappa})$ (see Fig. SN4a), and hence is favored also when $\kappa + \bar{\kappa} > 0$ (see Fig. SN4b, dashed line). This criteria hence determines the conditions for *trans* fusion and stability of a system of spherical lobules. On the other hand, high genus shapes are favored when $\bar{\kappa} > 0$ and *cis* fusion lowers bending energy by $\Delta E_b \simeq -4\pi\bar{\kappa}$ (see Fig. SN4a).

Using these conditions we propose a simple two dimensional state diagram (see Fig. SN4b), where we consider a system of spheres as initial state. Here three morphological states are found. We can further simplify this state diagram for $\kappa > 0$ into a one-dimensional state diagram as a function of the reduced Gaussian rigidity $\bar{\kappa}/\kappa$ (see Main text and Fig. 3b).

Resultant morphologies can then be interpreted by studying the variation of only one parameter, $\bar{\kappa}/\kappa$. High genus structures lie in region III, where $\bar{\kappa}/\kappa > 0$. A system of spheres and tubes, formed via *trans* fusion, lie in region II for which $\bar{\kappa}/\kappa > -1$.

More generally for any system of $N$ lobules and $g$ handles ($g - 1$ passages)

$$E_b = 4\pi N(\kappa s_l + \bar{\kappa}) - 4\pi\bar{\kappa} g \quad , \tag{30}$$

where $s_l = (\int H^2 dA)/N$ is a number that captures the average shape energy of the lobules. Note that for a system of spheres $s_l = 1$ and it is shown by Willmore that $s_l \geq 1$ (*19*). In that case the condition for *trans* fusion being favored yields a correction $\bar{\kappa}/\kappa > -s_l$. This modified criteria is naturally satisfied when $\bar{\kappa}/\kappa > -1$ as $s_l \geq 1$, hence for simplicity we limit our discussion to $s_l \sim 1$.

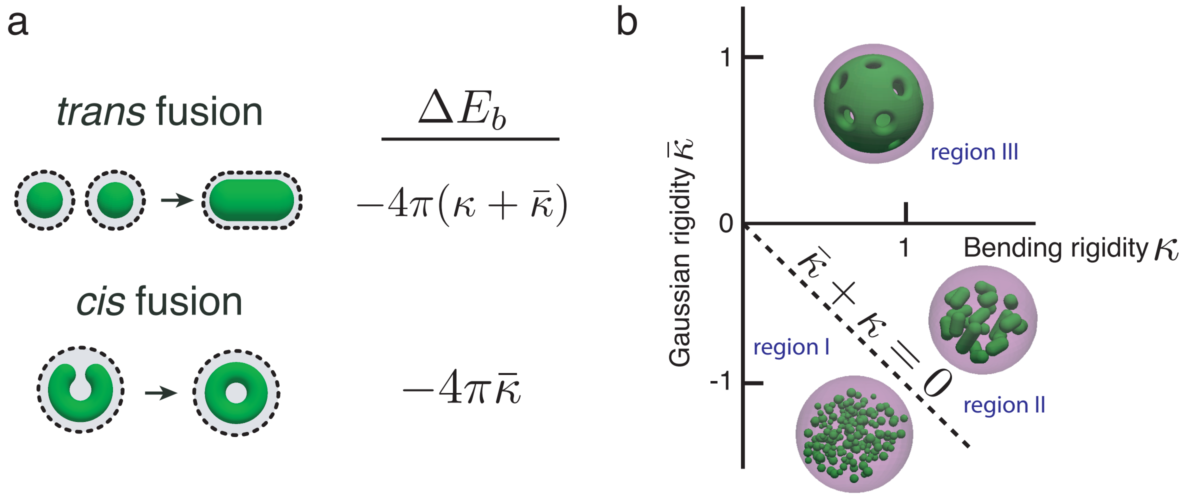


Fig. SN4: **a**, Changes in bending energy $\Delta E_b$ is shown for examples of *trans* (top) and *cis* (bottom) fusion. **b**, A state diagram representing three regions. In region I (below dashed line), fusions are disfavored, while in region II only *trans* fusion is favored. Both *trans* and *cis* fusion are favored in region III giving rise to structures with many passages. Insets show example systems with representative topology.

#### 2.2 Morphogenesis guided by *trans* fusion

In this section we discuss how successive *trans* fusion of almost spherical lobules can give rise to tubular shapes and as a result drive the morphological trajectory of neuroepithelial organoids. To discuss this we use the idealized geometric model of spherocylinder, which in the limit of vanishing aspect ratio ($a = 0$) is equivalent to a sphere.

Consider the *trans* fusion of two spheres of equal radii $R_0$, where the total surface area and volume remains unchanged by the fusion event. This geometric constraint, then determines the resultant geometry of the final shape. The area and the volume of the final shape is then given by

$$A = 8\pi R_0^2 \quad , \quad V = \frac{8}{3}\pi R_0^3 \quad . \tag{31}$$

The reduced volume $v$ for this final shape is thus $v = 1/\sqrt{2} \simeq 0.707$. For fluid surfaces of almost spherical geometry, prolate structures are found to be stable (*20*).

Such prolate shapes are well represented by the spherocylinder family described in the previous sections. The shape of a spherocylinder of radius $R$ and length $L$ formed by coaslescence of $n$ identical spheres of radius $R_0$ is characterized then by its aspect ratio $a$, which is a function of $n$. The aspect ratio $a(n)$ can be obtained by solving the area and volume constraint given by

$$4\pi R^2 + 2\pi RL = n4\pi R_0^2 \quad , \quad \frac{4}{3}\pi R^3 + \pi R^2 L = n\frac{4}{3}\pi R_0^3 \quad . \tag{32}$$

The exact solution of $a(n)$ can be obtained (see Fig. SN5a, blue solid line). To a good numerical approximation $a(n) \sim (2/\sqrt{\pi})\sqrt{n-1} + (13/3)(n-1)$ (see Fig. SN5a , yellow dashed line).

We find that the aspect ratio $a$ to be a monotonic function of $n$, hence implying how progressive trans fusion of spherical lobules can give rise to elongate tubes with increasing aspect ratio. This will result in decreasing reduced volume $v$ and increasing reduced curvature $m$ (see Fig. SN5b,c), depicting a trajectory in $(v, m)$ space similar to observed morphogenetic trajectory of neuroepithelial organoids (see Fig. 2a).

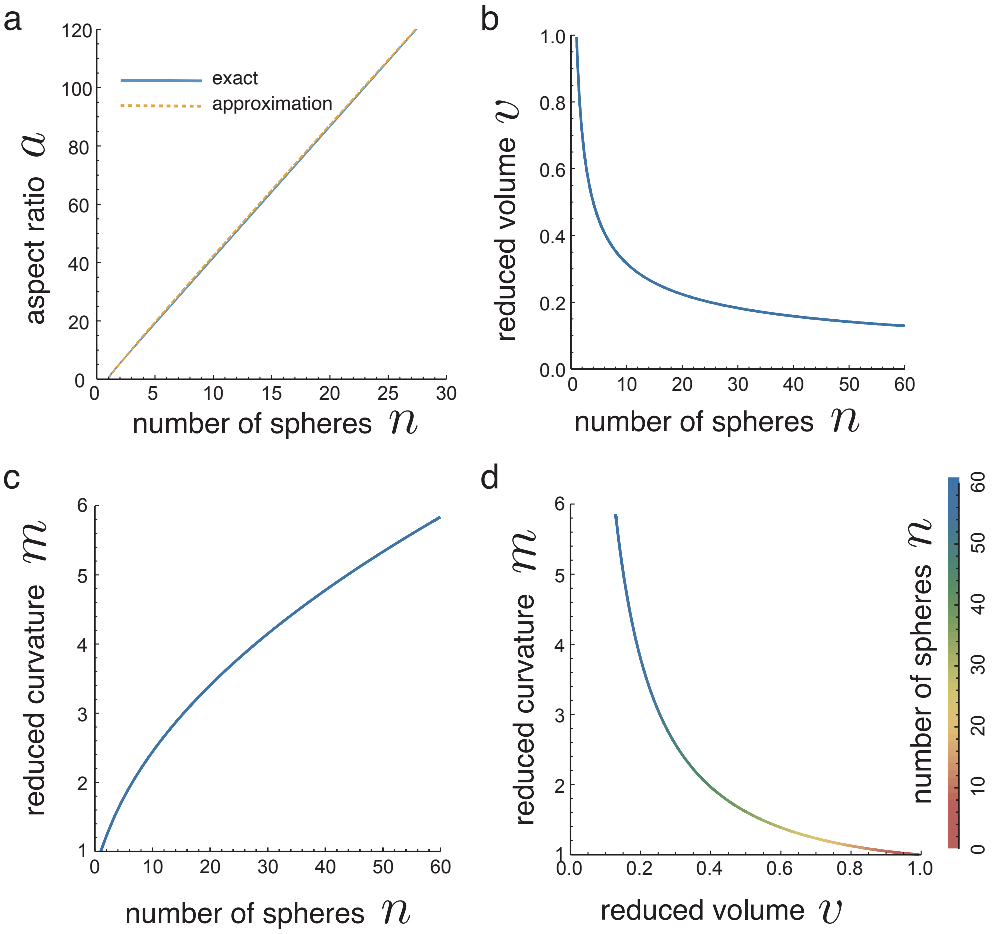


Figure SN5: **a**, Aspect ratio $a$ of a spherocylinder formed by coalescence of $n$ identical spheres is shown as a function of $n$. The approximate solution (yellow dashed line, see text) lies closely to the exact solution (blue solid line) for $a(n)$. **b**,**c**, Reduced volume $v$ and reduced curvature $m$ of the said spherocylinder is shown as a function of $n$. **d**, Parametric curve depicting morphogenetic trajectory of a spherocylinder in $(v, m)$ space parametrized by $n$.

5. Pauli Virtanen, Ralf Gommers, Travis E Oliphant, Matt Haberland, Tyler Reddy, David Cournapeau, Evgeni Burovski, Pearu Peterson, Warren Weckesser, Jonathan Bright, Stéfan J van der Walt, Matthew Brett, Joshua Wilson, K Jarrod Millman, Nikolay Mayorov, Andrew R J Nelson, Eric Jones, Robert Kern, Eric Larson, C J Carey, İlhan Polat, Yu Feng, Eric W Moore, Jake VanderPlas, Denis Laxalde, Josef Perktold, Robert Cimrman, Ian Henriksen, E A Quintero, Charles R Harris, Anne M Archibald, Antônio H Ribeiro, Fabian Pedregosa, Paul van Mulbregt, and SciPy 1.0 Contributors. SciPy 1.0: fundamental algorithms for scientific computing in Python. *Nature methods*, 17(3):261–272, March 2020.

6. . The pandas development team. pandas-dev/pandas: Pandas, February 2020.

16. Jakob Steiner. Über parallele flächen. *Monatsber. Preuss. Akad. Wiss*, 2:114–118, 1840.

17. Frank Jülicher. The morphology of vesicles of higher topological genus: conformal degeneracy and conformal modes. *Journal de physique II*, 6(12):1797–1824, 1996.

18. NJA Sloane, R Hardin, and Smith WD. Tables of Spherical Codes. NeilSloane.com/packings.

19. Thomas Willmore. *Total curvature in Riemannian geometry.* Ellis Horwood, 1982.

20. Udo Seifert, Karin Berndl, and Reinhard Lipowsky. Shape transformations of vesicles: Phase diagram for spontaneous-curvature and bilayer-coupling models. *Physical review A*, 44(2):1182, 1991.

#### Table Caption

- **Table S1.** This table lists the genes that were differentially regulated in response to retinoic acid (RA) treatment as identified from the RNA sequencing of organoids. Columns represent (1) Gene Symbol, (2) $\log_2$ fold change of transcript, RA-treated over untreated organoids, (3) $-\log_{10}(p_{adj})$, and (4) $p_{adj}$, where $p_{adj}$ is the adjusted p-value of statistical significance.

#### Movie Captions

- **Movie S1-2.** Surface representations of an untreated organoid (movie S1) and retinoic acid (RA)-treated organoid (movie S2).

  Same organoids as in Fig. 1b. Organoid outer boundary (magenta) and apical surfaces (green) were reconstructed from Day 4 organoids stained for DNA and anti-PODXL.

- **Movie S3-4.** Live imaging of an untreated organoid (Movie S3) and retinoic acid (RA)-treated organoid (Movie S4) undergoing morphogenesis from Day 2 to 4. Maximum intensity projections of SiR-actin volumetric images are shown as movies.

- **Movie S5-6.** Surface rendering of untreated organoid morphogenesis (Movie S5, corresponding to SiR-actin Movie S3) and RA-treated organoid morphogenesis (Movie S6, corresponding to SiR-actin Movie S4). Organoid outer boundary (magenta, transparent) and apical surfaces (green).

- **Movie S7-8.** The mean curvature $H$ of apical surfaces is displayed via a blue-red (low-high) colormap during the Day 2-4 morphogenesis of untreated (Movie S7) and RA-treated (movie S8) organoids.

- **Movie S9-10.** The Gaussian curvature $K$ of apical surfaces is displayed via a purple-green (low-high) colormap during the Day 2-4 morphogenesis of untreated (Movie S9) and RA-treated (Movie S10) organoids.

- **Movie S11-12.** Surface representations of an organoid treated with 2 $\mu$M HA130 (Movie S11) and an organoid treated with RA and 2 $\mu$M HA130 (Movie S12). Same organoids as in Fig. 3d. Organoid outer boundary (magenta) and apical surfaces (green) were reconstructed from Day 4 organoids stained for DNA and anti-PODXL. For Movies S3 to S10, time indicates hours after the Day 2 time point when new media with or without RA was added to the culture (see Fig. 2a, experimental timeline).
